# Long-term association of pregnancy and maternal brain structure: the Rotterdam Study

**DOI:** 10.1101/2021.02.19.432038

**Authors:** Jurate Aleknaviciute, Tavia E. Evans, Elif Aribas, Merel W. de Vries, Eric A.P. Steegers, M. Arfan Ikram, Henning Tiemeier, Maryam Kavousi, Meike W. Vernooij, Steven A. Kushner

## Abstract

The peripartum period is the highest risk interval for the onset or exacerbation of psychiatric illness in women’s lives. Notably, pregnancy and childbirth have been associated with short-term structural and functional changes in the maternal human brain. Yet the long-term effects of parity on maternal brain structure remain unknown. Therefore, we utilized a large population-based cohort to examine the association between parity and brain structure. In total, 2,835 women (mean age 65.2 years; all free from dementia, stroke, and cortical brain infarcts) from the Rotterdam Study underwent magnetic resonance imaging (1.5 T) between 2005 and 2015. Associations of parity with global and lobar brain tissue volumes, white matter microstructure, and markers of vascular brain disease were examined using regression models. We found that parity was associated with a larger global gray matter volume (β= 0.14, 95% CI = 0.09-0.19), a finding that persisted following adjustment for sociodemographic factors. A non-significant dose-dependent relationship was observed between a higher number of childbirths and larger gray matter volume. The gray matter volume association with parity was globally proportional across lobes. No associations were found regarding white matter volume or integrity, nor with markers of cerebral small vessel disease. The current findings indicate that pregnancy and childbirth are associated with robust long-term changes in brain structure involving larger global gray matter volume that persists for decades. Taken together, these data provide novel insight into the impact of motherhood on the human brain.

## INTRODUCTION

Pregnancy and childbirth are remarkable life-changing events, from personal, social, and biological perspectives. Motherhood is an extensive adaptation, altering behavior, motivation, and emotion in the service of offspring care. Such peripartum changes in brain function have been postulated as homeostatic mechanisms to mitigate the substantially elevated risk for the onset or exacerbation of psychiatric disorders in the postpartum period [1–5].

Apart from functional changes, there is growing evidence that the maternal brain exhibits considerable structural plasticity in association with pregnancy and parturition [6–8]. Studies using animal models provide evidence that pregnancy and parturition induce profound neurobiological changes on the maternal brain in rodents [9–12]. However, few brain imaging studies have been performed to examine the structural brain changes that occur in women during pregnancy and the postpartum period. The few that do exist concluded that pregnancy is associated with a reduction of gray matter volume [13,14]. Hoekzema and collegues [14] found that the observed postpartum reductions in specific regions of cortical gray matter remained evident two years after childbirth, suggesting that pregnancy can exert enduring structural changes on the human maternal brain. However, a study by Oatridge [13] following a small sample of healthy pregnant women serving as a control group for women with pre-eclampsia throughout pregnancy and the early postpartum period showed that global brain volume decreased during pregnancy with a nadir around the time of delivery, followed by a return to pre-pregnancy global brain volume within six months postpartum. Additional support for increasing brain size during the early postpartum period comes from a longitudinal within-subject analysis comparing images acquired 2-4 weeks postpartum with those acquired 3-4 months postpartum, which demonstrated increases of cortical gray matter volume in multiple brain regions [15]. A more recent study applied a machine learning algorithm to a longitudinal within-subject brain imaging dataset to estimate brain age in the first 2 days postpartum and again at 4-6 weeks postpartum, which revealed a “younger” brain age at 4-6 weeks postpartum [16].

Taken together, the currently available data suggest a robust morphological plasticity of the human maternal brain during pregnancy and the early postpartum period. However due to relatively short and differential postpartum periods in earlier studies, the nature of the relationship between pregnancy and childbirth with brain structure remains unresolved. Considering mounting evidence of the effects of neuroendocrine physiology on brain structure and function [17, 18], it is plausible that pregnancy and childbirth might present a critical yet understudied factor that is crucial for understanding the brain processes over the life course. Therefore, we utilized data from the population-based Rotterdam Study cohort to investigate the long-term association of parity with global brain structure and vascular integrity.

## METHODS AND MATERIALS

### Setting and participants

The Rotterdam Study, a prospective population-based cohort study, includes approximately 18,000 participants aged 40 years and older living in Ommoord, a suburb of Rotterdam [17]. Since its inception in 1990, participants have undergone follow-up visits to the research center at 3 to 6 year intervals. At interview female participants were asked about their reproductive status including parturition history. A dedicated magnetic resonance (MR) imaging scanner, with a fixed brain MRI protocol, was added in 2005 to the core study protocol (the Rotterdam Scan Study) [18]. Since brain MRI was introduced, more than 12,000 scans in 5,913 individuals have been acquired and processed, of which 3,257 are women. Of this sample, 3,197 women have imaging data and information on childbirth, of which 309 had dementia or a stroke at scan date and an additional 54 had cortical infarcts on brain MRI. The final cohort included 2,834 stroke-, dementia and cortical infarct-free women.

The Rotterdam Study has been approved by the Medical Ethics Committee of the Erasmus MC (registration number MEC 02.1015) and by the Dutch Ministry of Health, Welfare, and Sport (Population Screening Act WBO, license number 1071272-159521-PG). The Rotterdam Study has been entered into the Netherlands National Trial Register (NTR; www.trialregister.nl) and into the WHO International Clinical Trials Registry Platform (ICTRP; www.who.int/ictrp/network/primary/en/) under shared catalog number NTR6831. All participants provided written informed consent to participate in the study and to have their information obtained information from their treating physicians.

### Sociodemographic, pregnancy and parity characteristics

At study entry, women were asked by trained interviewers regarding data on parity and maternal age at first childbirth, as well as sociodemographic factors. We defined parity as the number of pregnancies with a gestational age at delivery of over 24 weeks. Level of education was defined categorically as either primary education, lower general and vocational education, intermediate/higher general and intermediate vocational education, or higher vocational education/ university. Smoking history was categorized as current, former, or never. Marital history was categorized as never- or ever-married. Information on body mass index (BMI) was obtained through interview and physical examination at a center visit closest in date to the MRI acquisition [19] and was calculated as weight (kg)/height^2^(m).

From January 2012 onwards, an additional questionnaire was added to the study protocol regarding having experienced complications during pregnancies. This question included history of any clinically diagnosed pregnancy complication, including gestational diabetes, high blood pressure during pregnancy, and pre-eclampsia, eclampsia or Hemolysis, elevated liver enzymes, and low platelet (HELLP) syndrome. This questionnaire was only available in a selected sample (894 of the women who had pregnancy information) as women who in the previous visits before 2012 had indicated they have already gone through menopause were not asked to participate in the reproductive questionnaire, including the new set of questions on pregnancy complications.

### MRI acquisition

MR images were acquired on a 1.5 tesla MRI scanner with an 8-channel head coil (GE signa Excite, General Electric Healthcare, Milwaukee, USA). The sequences acquired included a T1-weighted (T1w), proton density-weighted (PDw) and a fluid inversion recovery (FLAIR), all of which were used for automated brain tissue segmentation. Additionally, a T2^*^-weighted sequence was acquired. Full information on the scan parameters has been described previously (20). Diffusion imaging was performed via an echo planar imaging (EPI) readout with gradients (b=1,000 s/mm^2^) applied in 25 directions. Diffusion data were processed using a standardized pipeline, as described previously [20].

Tissue segmentation was performed using an automated processing algorithm based on a k-nearest-neighbour-classifier on the T1w and PDw scans complemented with FLAIR-intensity based white matter hyperintensity detection. The images were segmented into gray matter, cerebrospinal fluid, normal-appearing white matter (NAWM) and white matter hyperintensities [21,22]. Tissue segmentation results for white matter hyperintensities and NAWM were combined with the diffusion maps to extract global measures of fractional anisotropy and mean diffusivity within NAWM. Intracranial volume (ICV) was computed by the sum of all brain tissue classes and cerebrospinal fluid. Segmentation of the lobar regions, temporal-, frontal-, parietal- and occipital lobes, were carried out with left and right hemisphere volumes averaged for the analysis [23]. All segmentations were visually inspected by trained raters and corrected manually when needed [20, 23].

### Infarct and microbleed rating

Scans were visually inspected for the presence of lacunar infarcts and cerebral microbleeds by trained research physicians. Lacunar infarcts were defined on T2-weighted images as focal hyperintensities equal or larger than 3mm, but smaller than 15mm in size, and exhibiting the same characteristics as cerebrospinal fluid on all sequences - a hyperintense rim on the FLAIR sequence when located supratentorially with no involvement of the cortical gray matter. Microbleeds were defined as focal, small, round to ovoid areas of signal loss on T2^*^-weighted images, as previously defined [20].

### Statistical analysis

Regression analysis was performed with nulliparous/parous and parity groups (primiparous, 2-3 term pregnancies, and 4+ term pregnancies), with nulliparous women (parity = 0) as a reference in relation to imaging markers. Additionally, a stratified subgroup analysis was performed between 3 groups: nulliparous, parous women without complications during pregnancy, and parous women with complications during pregnancy. White matter hyperintensity volume was log-transformed due to a skewed distribution All variables except lacunar infarcts and microbleeds were *z*-transformed for comparison. Model I adjusted for age, and ICV. Model II further adjusted for education, marital status, smoking and body mass index. Analyses with DTI parameters as outcome were further adjusted for white matter volume and log-transformed white matter lesion volume in both models I and II.

#### Sensitivity analyses

Considering the suggested independent relationships of pre-eclampsia, menopause and hormone replacement therapy (HRT) with brain structure and function [17, 18, 24, 25] we conducted two sensitivity analyses to examine whether the results of the potential effect of pregnancy on brain structure were altered by pregnancy-related complications and menopause and hormone replacement therapy. The first analysis was restricted to a subset of 894 women for whom the additional information on pregnancy complications was available. Of these, 230 women reported having experienced complications such as including gestational diabetes, high blood pressure during pregnancy, and pre-eclampsia, eclampsia or HELLP syndrome, during pregnancy.

The second analysis was conducted in a subset of 1529 women for whom menopause and HRT data were available. Details of menopause and use of hormone replacement therapy were obtained by self-report during a home interview by trained interviewers and verified through general practitioner records. Women in this subset were classified based on menopause status and use of HRT (premenopausal [n=385], post-menopausal [n=830], post-menopausal + HRT [n=314]).

## RESULTS

In total, 2,834 women were included in the analyses. Of these, 441 were nulliparous (parity = 0) and 2,393 were parous (parity >=1). Further categorizing the group of parous women, 474 women were primiparous, 1,697 women had 2-3 pregnancies, and 222 women had 4 or more pregnancies. The sample characteristics classified by parity are shown in Table 1.

**Table 1.** Group characteristics

|  | Nulliparous | Primiparous | Multiparous<br>(parity = 2-3) | Multiparous<br>(parity ≥ 4) |
| --- | --- | --- | --- | --- |
| Number, n | 441 | 474 | 1,697 | 222 |
| **characteristics** |  |  |  |  |
| Age at time of MRI, years | 64.45 (11.51) | 62.93 (9.81) | 64.20 (10.42) | 69.08 (12.25) |
| Age at first child, years | - | 27.56 (5.4) | 25.23 (4) | 23.96 (3.51) |
| Married, ever. % (n) | 72 (288) | 97 (415) | 99 (1555) | 99 (199) |
| Education, n |  |  |  |  |
| 0 | 35 | 57 | 148 | 30 |
| 1 | 168 | 245 | 854 | 96 |
| 2 | 131 | 92 | 405 | 69 |
| 3 | 105 | 77 | 284 | 28 |
| Body mass index, kg/m<sup>2</sup> | 26.64 (4.83) | 27.03 (4.70) | 27.56 (4.52) | 28.05 (4.73) |
| Smoking, n |  |  |  |  |
| Never | 146 | 177 | 684 | 114 |
| Past smoking | 208 | 191 | 743 | 76 |
| Current smoking | 87 | 103 | 264 | 29 |
| **Brain volumes** |  |  |  |  |
| Total brain volume, ml | 905.99 (81.46) | 902.65 (83.42) | 900.88 (81.86) | 883.45 (94.44) |
| Gray matter volume, ml | 505.65 (45.92) | 506.69 (43.09) | 507.93 (44.91) | 500.07 (50.20) |
| White matter volume, ml | 393.22 (50.69) | 389.59 (52.46) | 387.05 (52.91) | 375.15 (10.40) |
| Frontal lobe volume, ml | 85.48 (8.69) | 85.65 (8.43) | 85.86 (8.60) | 84.27 (9.87) |
| Temporal lobe volume, ml | 58.81 (5.19) | 59.07 (5.05) | 59.00 (5.11) | 57.75 (5.62) |
| Occipital lobe volume, ml | 31.66 (3.61) | 31.87 (3.25) | 31.93 (3.41) | 31.27 (3.52) |
| Parietal lobe volume, ml | 49.92 (5.30) | 49.89 (4.85) | 50.16 (5.15) | 49.63 (5.60) |
| **Brain Microstructure** |  |  |  |  |
| Fractional anisotropy | 0.34 (0.02) | 0.34 (0.02) | 0.34 (0.01) | 0.34 (0.02) |
| Mean diffusivity, 10<sup>-3</sup> mm<sup>2</sup>/s | 0.75 (0.03) | 0.74 (0.03) | 0.74 (0.03) | 0.76 (0.04) |
| **Markers of cerebral small vessel disease** |  |  |  |  |
| White matter hyperintensity volume, ml | 7.12 (13.18) | 6.37 (11.59) | 5.90 (8.47) | 8.23 (10.40) |
| Lacunar infarct (Y/N) | 4% | 6% | 6% | 7 % |
| Microbleed (Y/N) | 20% | 17% | 17% | 29% |

Values are reported as means (standard deviations) unless stated otherwise. Missing values were present in education (65), marital status (243), smoking (16). Education: (0) primary education, (1) Lower general and vocational education, (2) Intermediate/higher general and intermediate vocational education, (3) higher vocational education/ university.

With respect to brain tissue volumes, we found that parity was associated with a larger global gray matter volume [adjusted mean difference (β)= 0.14, 95% confidence intervals (CI) = 0.09;0.19] (Tables 2–3). This association persisted following adjustment for smoking, BMI, education, and history of marital status (β = 0.10, 95% CI = 0.04;0.17). We found no differences in white matter volume associated with parity. The relationship between gray matter volume and parity was consistent across temporal, frontal, occipital and parietal regions (Supplementary Table 1, Model I). These relationships were attenuated after adjustment for sociodemographic factors (Supplementary Table 1, Model II). No disproportionate lobar changes were found.

**Table 2.** Relationship between parity (parous/nulliparous) and structural brain imaging markers

|  | Model I | Model II |
| --- | --- | --- |
|  | Beta (95% CI) | Beta (95% CI) |
| Total brain volume | **0.08 (0.04;0.12)** | **0.07 (0.03;0.12)** |
| Gray matter volume | **0.14 (0.09;0.19)** | **0.11 (0.04;0.17)** |
| White matter volume | 0.00 (-0.05;0.06) | 0.02 (-0.04;0.09) |
| Fractional anisotropy | 0.05 (-0.04;0.15) | 0.03 (-0.08;0.14) |
| Mean Diffusivity | **-0.07 (-0.14;0.00)** | -0.06 (-0.14;0.02) |
| White matter hyperintensity volume | -0.01 (-0.09;0.06) | -0.00 (-0.08;0.08) |
| Lacunar Infarct | 0.02 (-0.01;0.04) | 0.01 (-0.01;0.04) |
| Microbleed | -0.01 (-0.05;0.03) | 0.01 (-0.04;0.05) |

White matter lesion volume was log-transformed due to a skewed distribution. All variables except infarct and microbleeds were z-transformed to allow for comparison. Bold indicates $p < 0.05$ .
Model I was adjusted for age and intracranial volume. Model II further adjusted for education, body mass index, smoking and marital history. DTI analysis further adjusted for white matter volume and white matter lesion volume in both models I and II.

**Table 3.** Relationship between the parity and structural brain imaging markers

| Nulliparous as reference | Primiparous | Multiparous (parity =2-3) | Multiparous (parity ≥4) |
| --- | --- | --- | --- |
|  | Beta (95% CI) | Beta (95% CI) | Beta (95% CI) |
| **Model I** |  |  |  |
| Total brain volume | **0.07(0.02;0.12)** | **0.09(0.05;0.13)** | **0.07 (0.01;0.13)** |
| Gray matter volume | **0.10 (0.03;0.17)** | **0.15 (0.09;0.21)** | **0.15 (0.06;0.24)** |
| White matter volume | 0.01 (-0.06;0.08) | 0 (-0.06;0.06) | -0.02 (-0.11;0.07) |
| Fractional anisotropy | 0 (-0.12;0.12) | 0.08 (-0.01;0.17) | -0.01 (-0.16;0.13) |
| Mean Diffusivity | **-0.08 (-0.17;0.00)** | **-0.08 (-0.15;-0.01)** | 0.00 (-0.10;0.11) |
| White matter hyperintensity volume | 0.03 (-0.06;0.12) | -0.03 (-0.1;0.05) | -0.01 (-0.12;0.11) |
| Lacunar Infarct | 0.02 (-0.01;0.05) | 0.02 (-0.01;0.04) | 0.02 (-0.02;0.05) |
| Microbleed | -0.01 (-0.06;0.04) | -0.02 (-0.06;0.02) | 0.05 (-0.01;0.12) |
| **Model II** |  |  |  |
| Total brain volume | **0.06(0.01;0.11)** | **0.07 (0.03;0.12)** | **0.09 (0.02;0.15)** |
| Gray matter volume | **0.08 (0.00;0.16)** | **0.11 (0.05;0.18)** | **0.13 (0.03;0.22)** |
| White matter volume | 0.02 (-0.06;0.10) | 0.02 (-0.05;0.09) | 0.02 (-0.07;0.12) |
| Fractional anisotropy | -0.04 (-0.17;0.09) | 0.06 (-0.05;0.17) | -0.03 (-0.19;0.13) |
| Mean Diffusivity | -0.07 (-0.17;0.03) | -0.06 (-0.15;0.02) | 0.01 (-0.11;0.13) |
| White matter hyperintensity volume | 0.02 (-0.08;0.12) | -0.01 (-0.09;0.08) | 0.02 (-0.11;0.14) |
| Lacunar Infarct | 0.02 (-0.01;0.06) | 0.01 (-0.02;0.04) | 0.01 (-0.03;0.05) |
| Microbleed | 0.00 (-0.06;0.06) | 0.00 (-0.05;0.05) | **0.08(0.01;0.14)** |

White matter lesion volume was log-transformed due to a skewed distribution All variables except infarct and microbleeds were z-transformed to allow for comparison. Bold indicates $p < 0.05$ .
Model I was adjusted for age and intracranial volume. Model II further adjusted for education, body mass index, smoking and marital history. DTI analysis further adjusted for white matter volume and white matter lesion volume in both models I and II.

For analyses involving microstructural outcomes, mean diffusivity was lower in parous women (β = −0.07, 95% CI = −0.14;0.00). No relationships were observed between parity and fractional anisotropy in normal-appearing white matter (NAWM). There was also no relationship between parity and markers of cerebral small vessel disease, with the exception of an increase of microbleeds observed in multiparous (parity ≥ 4) women compared to nulliparous women [β = 0.07, 95% CI = 0.01;0.13] (Table 3, Model II).

We next examined whether the association between parity and brain structure might be differentially influenced by parity. Nulliparous women were considered as the reference group. The results demonstrated a non-significant dose-dependent trend in the association between parity and total gray matter volume (Table 3, Model I in reference to nulliparous women; primiparous [β = 0.10, 95% CI = 0.03;0.17], multiparous women (parity: 2-3) [β = 0.15, 95% CI = 0.09;0.21], multiparous women (parity ≥ 4) [β = 0.15, 95% CI = 0.06;0.24]). However, these associations were attenuated after adjustment for sociodemographic factors (Table 3, Model II; primiparous [β = 0.07, 95% CI = −0.00;0.15], multiparous (parity = 2-3) [β = 0.12, 95% CI = 0.05;0.18], multiparous (parity ≥4) [β = 0.12, 95% CI = 0.02;0.22]). No other brain imaging marker studied exhibited a dose-dependent relationship with parity (Table 3).

### Sensitivity analysis

In total, 894 women had information on pregnancy-related complications. Of these, 664 women reported no complications, and 230 women reported a history of complications during pregnancy. The subsample characteristics are shown in Table 4, stratified by pregnancy complications (parous without pregnancy complications, parous with pregnancy complications).

**Table 4.**
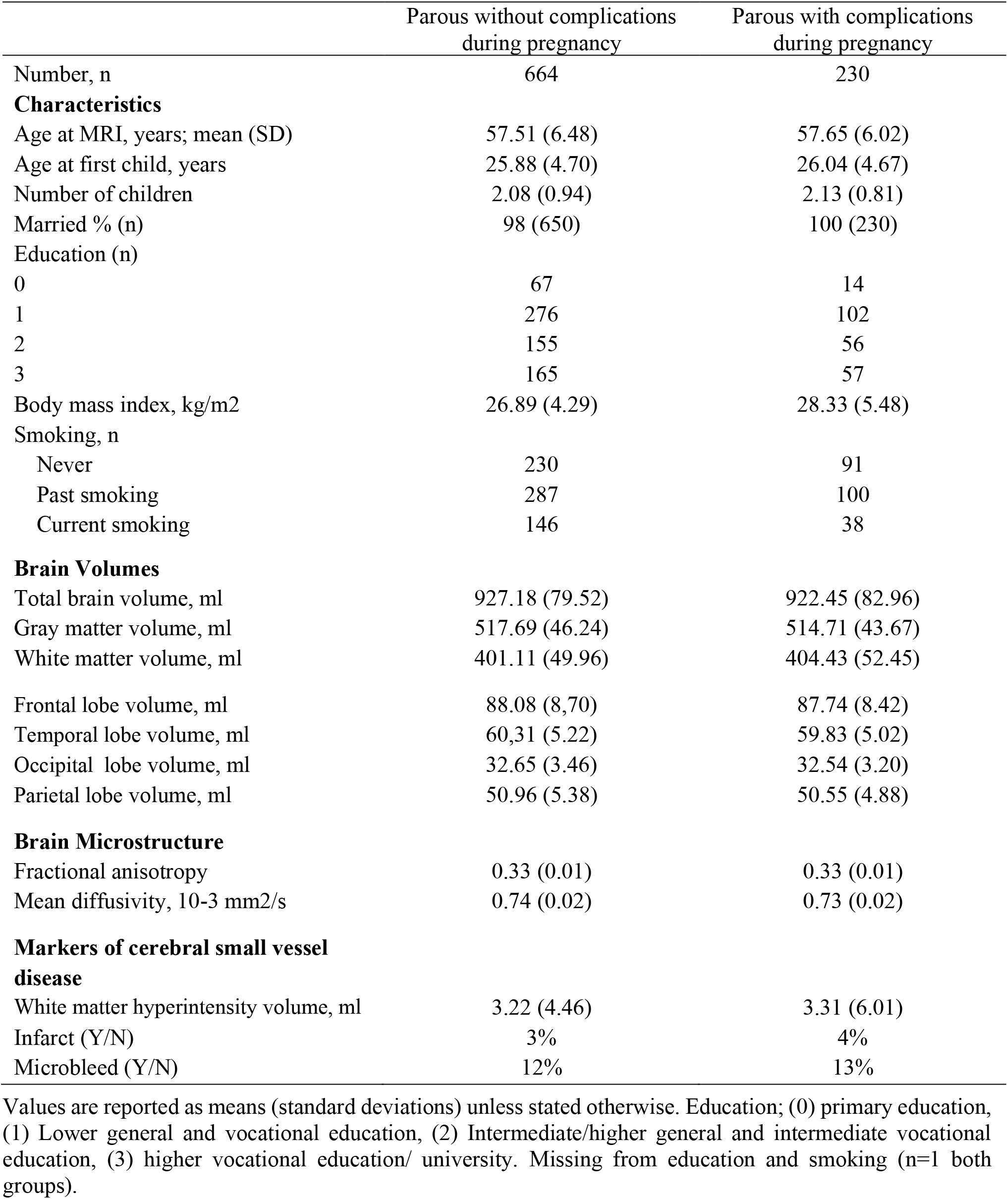
Pregnancy-related complications: group characteristics

|  | Parous without complications<br>during pregnancy | Parous with complications<br>during pregnancy |
| --- | --- | --- |
| Number, n | 664 | 230 |
| **Characteristics** |  |  |
| Age at MRI, years; mean (SD) | 57.51 (6.48) | 57.65 (6.02) |
| Age at first child, years | 25.88 (4.70) | 26.04 (4.67) |
| Number of children | 2.08 (0.94) | 2.13 (0.81) |
| Married % (n) | 98 (650) | 100 (230) |
| Education (n) |  |  |
| 0 | 67 | 14 |
| 1 | 276 | 102 |
| 2 | 155 | 56 |
| 3 | 165 | 57 |
| Body mass index, kg/m<sup>2</sup> | 26.89 (4.29) | 28.33 (5.48) |
| Smoking, n |  |  |
| Never | 230 | 91 |
| Past smoking | 287 | 100 |
| Current smoking | 146 | 38 |
| **Brain Volumes** |  |  |
| Total brain volume, ml | 927.18 (79.52) | 922.45 (82.96) |
| Gray matter volume, ml | 517.69 (46.24) | 514.71 (43.67) |
| White matter volume, ml | 401.11 (49.96) | 404.43 (52.45) |
| Frontal lobe volume, ml | 88.08 (8.70) | 87.74 (8.42) |
| Temporal lobe volume, ml | 60.31 (5.22) | 59.83 (5.02) |
| Occipital lobe volume, ml | 32.65 (3.46) | 32.54 (3.20) |
| Parietal lobe volume, ml | 50.96 (5.38) | 50.55 (4.88) |
| **Brain Microstructure** |  |  |
| Fractional anisotropy | 0.33 (0.01) | 0.33 (0.01) |
| Mean diffusivity, 10<sup>-3</sup> mm<sup>2</sup>/s | 0.74 (0.02) | 0.73 (0.02) |
| **Markers of cerebral small vessel disease** |  |  |
| White matter hyperintensity volume, ml | 3.22 (4.46) | 3.31 (6.01) |
| Infarct (Y/N) | 3% | 4% |
| Microbleed (Y/N) | 12% | 13% |

Values are reported as means (standard deviations) unless stated otherwise. Education; (0) primary education, (1) Lower general and vocational education, (2) Intermediate/higher general and intermediate vocational education, (3) higher vocational education/ university. Missing from education and smoking (n=1 both groups).

Evaluation of the subsample of women for whom information on pregnancy-related complications was available yielded differences exclusively in white matter between women with complications during pregnancy compared to parous women without pregnancy-related complications. Women with pregnancy-related complications exhibited larger white matter volumes (Table 5, Model I; β = 0.08, 95% CI = 0.01;0.16). This relationship attenuated after adjustment for smoking, BMI, and education (Table 5). Analysis of the relationship of parity with gray matter volume and MD showed similar effects as in the overall sample (Table 5, Model 1).

**Table 5.** Relationship between pregnancy complications and parity and structural brain imaging markers

|  | Parous with complications -<br>Parous without complications<br>during pregnancy (ref) | Parous without complications<br>during pregnancy -<br>Nulliparous (ref) | Parous with complications<br>during pregnancy -<br>Nulliparous (ref) |
| --- | --- | --- | --- |
|  | Beta (95% CI) | Beta (95% CI) | Beta (95% CI) |
| **Model I** |  |  |  |
| Total brain volume | -0.04(-0.01;0.08) | **0.11(0.07;0.16)** | **0.15(0.09;0.21)** |
| Gray matter volume | -0.02 (-0.11;-0.05) | **0.14 (0.07;0.21)** | **0.11 (0.02;0.20)** |
| White matter volume | **0.08 (0.01;0.16)** | 0.05 (-0.02;0.012) | **0.13 (0.05;0.22)** |
| Fractional anisotropy | -0.04 (-0.17;0.09) | 0.04 (-0.15;0.08) | -0.08 (-0.23;0.07) |
| Mean diffusivity | -0.02 (-0.11;0.06) | **-0.13 (-0.21;-0.04)** | **-0.15 (-0.26;-0.04)** |
| White matter<br>hyperintensity volume | 0.06 (-0.03;0.15) | **-0.09 (-0.18;0.00)** | -0.02 (-0.14;0.09) |
| Lacunar Infarct | 0.00 (-0.02;0.03) | 0.01 (-0.01;0.04) | 0.01 (-0.02;0.05) |
| Microbleed | 0.00 (-0.04;0.05) | -0.02 (-0.07;0.02) | -0.02 (-0.08;0.04) |
| **Model II** |  |  |  |
| Total brain volume | 0.04(-0.01;0.09) | **0.09(0.04;0.14)** | **0.12(0.06;0.19)** |
| Gray matter volume | -0.01 (-0.09;0.08) | **0.09 (0.01;0.17)** | 0.07 (-0.03;0.16) |
| White matter volume | 0.07 (-0.01;0.15) | -0.07 (-0.01;0.15) | **0.13 (0.04;0.23)** |
| Fractional anisotropy | -0.02 (-0.15;0.11) | -0.08 (-0.21;0.04) | -0.09 (-0.26;0.07) |
| Mean diffusivity | -0.03 (-0.12;0.06) | -0.08 (-0.18;0.00) | **-0.12 (-0.25;0.00)** |
| White matter<br>hyperintensity volume | 0.01 (-0.08;0.11) | -0.03 (-0.12;0.07) | 0.01 (-0.12;0.13) |
| Lacunar Infarct | 0.00 (-0.02;0.03) | 0.01 (-0.02;0.03) | 0.01 (-0.03;0.04) |
| Microbleed | 0.01 (-0.05;0.06) | -0.02 (-0.07;0.03) | 0.01 (-0.06;0.19) |

White matter lesion volume was log-transformed due to a skewed distribution. All variables except infarct and microbleeds were z-transformed to allow for comparison. Bold indicates $p < 0.05$ . Model I was adjusted for age and intracranial volume. Model II further adjusted for education, body mass index, smoking and marital history. DTI analysis further adjusted for white matter volume and white matter lesion volume in both models I and II. Ref; reference variable

A sensitivity analysis for the influence of menopause status and HRT on the relationship between parity and larger gray matter volume yielded no significant effects (β = 0.10, 95% CI = 0.02;0.17) (Supplementary Table 2).

## DISCUSSION

This population-based study demonstrated an association between parity and brain structure decades following pregnancy and childbirth. Specifically, we found that parity was associated with larger total gray matter volume later in life, a finding that persisted following adjustment for sociodemographic factors. The larger gray matter volume associated with parity appeared not to be driven by specific lobar brain regions, but rather was globally proportional across lobes. Moreover, we did not find evidence for an association between parity and markers of cerebral small vessel disease, which supports the theory of the adaptation of the brain and cerebral circulation during pregnancy to maintain brain homeostasis, despite substantial peripartum hormonal and cardiovascular changes [24].

A recently published study suggested that pregnancy-induced reductions in gray matter remained evident for at least two years after childbirth, implying a long-term reduction in brain tissue volume, primarily located in specific lobe regions [14]. Here we found that decades after pregnancy, gray matter volume is actually larger, a finding that remained robust following adjustment for age, BMI, smoking, education, and marital status [26, 27]. Moreover, the association of parity and gray matter volume persisted independent of pregnancy-related complications. Despite extensive reports on associations between pre-eclampsia and changes in cortical volumes [13, 30–32], this finding is in line with a prior study reporting no influence of preeclampsia on the association of parity and global gray matter volume in the early postpartum period [13].

Surprisingly, white matter volume was larger in women with pregnancy-related complications versus nulliparous women. A possible link underlying the association of larger white matter volume and hypertension-related complications might relate to a homeostatic compensation for chronic vascular insufficiency. Alternatively, however, the finding of larger white matter volumes in women with pregnancy-related complications might result from the small difference in age of the cohort sets, as the sub-sample with information on pregnancy-related complications was younger at the time of MRI acquisition compared to the main cohort. Another possibility is that the finding might reflect a survival bias, in which the participants of the study who experienced pregnancy-related complications were a disproportionately healthier subgroup. We acknowledge that exclusion of women who suffered from stroke, dementia, or cortical infarct might have introduced a selection bias. However, our decision to exclude those women was predicated on the reliability of the imaging markers.

There is a consensus that pregnancy, delivery, and puerperium expose women to a diversity of health changes that extend beyond direct obstetric complications of pregnancy. For example, emerging evidence suggests that women in the early postpartum period have a substantially increased risk of a first-onset or exacerbation of psychiatric disorders, cardiovascular, and autoimmune diseases [33–35]. However, investigations of the long-term risks associated with pregnancy and delivery have been inconclusive [36–39]. While some studies have argued that childbirth is associated with accelerated cellular aging due to higher levels of oxidative stress [36], other studies have contradicted these findings by demonstrating elongated telomeres suggestive of an attenuation of risk [37, 40]. Although no unifying biological explanation has emerged to explain these apparently contrasting findings, it remains a distinct possibility that pregnancy and childbirth have an enduring influence on the endocrine system, and consequently on brain structure, long after childbirth. Pregnancy and childbirth are accompanied by dramatic changes in the hormonal profile. Prolactin, androgens, and estrogens exhibit multiple orders of magnitude increases to support pregnancy, fetal growth, and delivery [34, 41, 42], which have been suggested to modulate several forms of brain plasticity, including changes in glial proliferation, neuronal morphology, and neurogenesis [43, 44].

Another potential explanation of enduring effects of pregnancy on brain structure is the bi-directional trafficking of maternal and fetal cells throughout gestation, which can acquire long-term residence in the human brain [45–51]. Fetal cells have been found at the sites of inflammation and linked to preeclampsia and multiple autoimmune diseases [49, 52]. Inflammation has also been associated with structural and functional brain changes [52]. Fetal cells are able to integrate into maternal brain circuity and express appropriate immunochemical markers for brain tissues [46, 48]. However, the extent to which fetal microchimerism is tolerated and whether dynamic changes occur over time remain unknown.

In parallel with underlying biological mechanisms of the observed association between parity and brain structure, the experience of parenting may also alter the brain, which is assumed to be necessary to support sensitive and responsive caregiving. A small number of studies have reported structural and functional brain changes associated with parenthood in humans [15, 54, 55] and it has been shown that the duration of motherhood is associated with greater neural activation to infant-specific cues [56, 57]. Another study found that foster mothers demonstrated an association between brain activity and caregiving behavior comparable to the associations observed in biological mothers [58]. Furthermore, a recent neuroimaging study in older adults found a positive association between the number of offspring and cortical thickness, in both fathers and mothers [59].

Our study has several limitations. Although data were sampled from a large, prospective, longitudinal population-based study allowing us to adjust for several covariates, we did not have information on infertility and gravidity which might have improved our ability to adjust for potential confounders. Furthermore, availability of time-varying cardiovascular risk factors might have been helpful to assess the mediating effects on the relationships between parity and brain volumes. Additionally, our sample consisted of a predominantly middle-class population of Caucasian descent, which may restrict the generalizability of our findings. Moreover, this study utilized a cross-sectional design, which precluded firm conclusions regarding the causality of the observed results. Information regarding pregnancy complications were available in a smaller subset of women from whom ~25% reported having experienced any pregnancy-related complications, which is larger than prior prevalence estimates of pregnancy-related complications [60–62]. As this questionnaire – at its introduction in the Rotterdam Study – was not asked from all women, the possibility of selection bias cannot be entirely ruled out. Moreover, the data was acquired using a self-report questionnaire. Considering no external validation data such as use of medication or treatment for pregnancy complications was available [63], recall bias cannot be excluded.

In addition, we acknowledge that we cannot distinguish between the effects of pregnancy, parity, and parenting on structural changes of the brain. Furthermore, it is possible that the observed effect of parity is a result of smaller brain volumes among nulliparous women, rather than larger gray matter volume in parous women. The important point in this context is that nulliparous and parous women might differ in several ways regarding their partnerships and unplanned pregnancies. Living in a relationship with a partner might have cognitive and social challenges that result in enduring changes of brain volume. Although we adjusted for the history of marital status, we cannot rule out residual confounding by factors such as unregistered partnerships. Hence, considering the study design and advanced age of the cohort, the findings may be interpreted in several ways. One possible interpretation is that over the life course, pregnancy and childbirth lead to an increase in global gray matter volume. Alternatively, pregnancy and childbirth may serve as a protective factor for subsequent age-related brain atrophy. Lastly, it might be that parenting creates an enriched social network that is protective against brain ageing.

In conclusion, the current findings indicate that parity is associated with a relatively larger global gray matter volume, decades following childbirth. Although the mechanism and physiological relevance of the morphological alterations remain unknown, these data provide novel insight into the long-term impact of motherhood on the human brain.

## Acknowledgements

The Rotterdam Study is supported by the Erasmus MC University Medical Center and Erasmus University Rotterdam; The Netherlands Organisation for Scientific Research (NWO); The Netherlands Organisation for Health Research and Development (ZonMw); the Research Institute for Diseases in the Elderly (RIDE); The Netherlands Genomics Initiative (NGI); the Ministry of Education, Culture and Science; the Ministry of Health, Welfare and Sports; the European Commission (DG XII); and the Municipality of Rotterdam. We gratefully acknowledge the participants within the Rotterdam Study and the general practitioners and pharmacists of the Ommoord district.

## SUPPLEMENT

**Supplementary Table 1.** Relationship between parity and gray matter volumes

|  | primiparous-<br>nulliparous | Multiparous<br>(parity =2-3)-<br>nulliparous | Multiparous<br>(parity ≥4)-<br>nulliparous |
| --- | --- | --- | --- |
|  | Beta (95% CI) | Beta (95% CI) | Beta (95% CI) |
| ***Model I*** |  |  |  |
| Total | **0.10 (0.03;0.17)** | **0.15 (0.09;0.21)** | **0.15 (0.06;0.24)** |
| Frontal lobe | **0.09 (0.01;0.16)** | **0.14 (0.08;0.20)** | **0.16 (0.07;0.25)** |
| Temporal lobe | **0.12 (0.04;0.19)** | **0.13 (0.07;0.19)** | 0.09 (-0.01;0.18) |
| Occipital lobe | **0.12 (0.03;0.21)** | **0.16 (0.09;0.23)** | **0.14 (0.03;0.25)** |
| Parietal lobe | **0.09 (0.01;0.17)** | **0.15 (0.08;0.21)** | **0.16 (0.07;0.26)** |
| ***Model II*** |  |  |  |
| Total | **0.08 (0.00;0.16)** | **0.11 (0.05;0.18)** | **0.13 (0.03;0.22)** |
| Frontal lobe | 0.07 (-0.01;0.15) | **0.11 (0.04;0.18)** | **0.14 (0.04;0.24)** |
| Temporal lobe | **0.10 (0.02;0.19)** | **0.10 (0.03;0.17)** | 0.07 (-0.03;0.18) |
| Occipital lobe | **0.10 (0.00;0.20)** | **0.11 (0.03;0.20)** | **0.13 (0.00;0.25)** |
| Parietal lobe | 0.05 (-0.03;0.14) | **0.11 (0.04;0.18)** | **0.14 (0.03;0.25)** |

White matter lesion volume was log-transformed due to a skewed distribution. All variables were $z$ -transformed to allow for comparison. Bold indicates $p < 0.05$ . Model I was adjusted for age and intracranial volume. Model II further adjusted for education, body mass index, smoking and marital history.

**Supplementary Table 2.** Relationship between the parity (parous/nulliparous) and structural brain imaging markers after adjusting for menopause status and hormone replacement therapy.

|  | Model I + Menopause/HRT<br>Beta (95% CI) |
| --- | --- |
| Total brain volume | **0.11 (0.06;0.16)** |
| Gray matter volume | **0.10 (0.02;0.17)** |
| White matter volume | **0.07 (0.00;0.14)** |
| Fractional anisotropy | 0.03 (-0.09;0.15) |
| Mean Diffusivity | -0.04 (-0.12;0.04) |
| White matter hyperintensity volume | -0.01 (-0.10;0.08) |
| Lacunar Infarct | 0.01(-0.02;0.03) |
| Microbleed | -0.02 (-0.06;0.03) |

Model I adjusts for age and ICV. Additionally, menopause and hormone replacement therapy (HRT) is adjusted as a factor variable coded as 0: no menopause at scan, 1: menopause and use of HRT, 2: menopause and no use of HRT. DTI analysis further adjusted for white matter volume and white matter lesion volume.

## REFERENCES

1. Brunton PJ, Russell JA. The expectant brain: Adapting for motherhood. Nat Rev Neurosci 2008; 9: 11–25.

2. Bergink V, Armangue T, Titulaer MJ, Markx S, Dalmau J, Kushner SA. Autoimmune Encephalitis in Postpartum Psychosis. Am J Psychiatry 2015; 172: 901–908.

3. Maguire J, Mody I. GABA(A)R plasticity during pregnancy: relevance to postpartum depression. Neuron 2008; 59: 207–213.

4. Putnam KT, Wilcox M, Robertson-Blackmore E, Sharkey K, Bergink V, Munk-Olsen T, et al. Clinical phenotypes of perinatal depression and time of symptom onset: analysis of data from an international consortium. Lancet Psychiatry 2017; 4: 477–485.

5. Meltzer-Brody S, Howard LM, Bergink V, Vigod S, Jones I, Munk-Olsen T, et al. Postpartum psychiatric disorders. Nat Rev Dis Primers 2018; 4:18022. doi: 10.1038/nrdp.2018.22.

6. Kim P, Strathearn L, Swain JE. The maternal brain and its plasticity in humans. Horm Behav 2016; 77: 113–123.

7. Hillerer KM, Jacobs VR, Fischer T, Aigner L. The maternal brain: An organ with peripartal plasticity. Neural Plast 2014; 2014: 574159. https://doi.org/10.1155/2014/574159

8. Leuner B, Sabihi S. The birth of new neurons in the maternal brain: Hormonal regulation and functional implications. Front Neuroendocrinol. 2016; 41: 99–113.

9. Dey S, Chamero P, Pru JK, Chien MS, Ibarra-Soria X, Spencer KR, et al. Cyclic Regulation of Sensory Perception by a Female Hormone Alters Behavior. Cell 2015; 16: 1334–13344.

10. Dulac C, O’Connel L, Wu Z. Neural control of maternal and paternal behaviors. Science 2014; 345: 765–770.

11. Kinsley CH, Lambert KG. Reproduction-induced neuroplasticity: Natural behavioural and neuronal alterations associated with the production and care of offspring. J Neuroendocrinol 2008; 20: 515–525.

12. Gatewood JD, Morgan MD, Eaton M, McNamara IM, Stevens LF, MacBeth AH, et al. Motherhood mitigates aging-related decrements in learning and memory and positively affects brain aging in the rat. Brain Res Bull 2005; 66: 91–98.

13. Oatridge A, Holdcroft A, Saeed N, Hajnal J V., Puri BK, Fusi L, Bydder GM (2002): Change in brain size during and after pregnancy: Study in healthy women and women with preeclampsia. Am J Neuroradiol. 23: 19–26.

14. Hoekzema E, Barba-Müller E, Pozzobon C, Picado M, Lucco F, García-García D, et al. Pregnancy leads to long-lasting changes in human brain structure. Nat Neurosci 2017; 20: 287–296.

15. Kim P, Leckman JF, Mayes LC, Feldman R, Wang X, Swain JE. The plasticity of human maternal brain: Longitudinal changes in brain anatomy during the early postpartum period. Behav Neurosci 2010; 124:695–700.

16. Luders E, Gingnell M, Poromaa IS, Engman J, Kurth F, Gaser C. Potential Brain Age Reversal after Pregnancy: Younger Brains at 4–6 Weeks Postpartum. Neuroscience 2018; 386: 309–314.

17. Kantarci K, Tosakulwong N, Lesnick TG, Zuk SM, Lowe VJ, Fields JA, et al. Brain structure and cognition 3 years after the end of an early menopausal hormone therapy trial. Neurology 2018; 90: e1404–e1412.

18. Kantarci K, Tosakulwong N, Lesnick TG, Zuk SM, Gunter JL, Senjem ML, et al. Changes in Brain Structure Three Years After the End of Menopausal Hormone Therapies in a Randomized Controlled Trial. Alzheimer’s Dement 2017; 13: P570.

19. Ikram MA, Brusselle GGO, Murad SD, van Duijn CM, Franco OH, Goedegebure A, et al. The Rotterdam Study: 2018 update on objectives, design and main results. Eur J Epidemiol 2017; 32: 807–850.

20. Ikram MA, van der Lugt A, Niessen WJ, Koudstaal PJ, Krestin GP, Hofman A, et al. The Rotterdam Scan Study: design update 2016 and main findings. Eur J Epidemiol 2015; 30: 1299–1315.

21. Boer de R, Vrooman HA, van der Lijn F, Vernooij MW, Ikram MA, van der Lugt A, et al. White matter lesion extension to automatic brain tissue segmentation on MRI. Neuroimage 2009; 45: 1151–1161.

22. Vrooman HA, Cocosco CA, van der Lijn F, Stokking R, Ikram MA, Vernooij MW, et al. Multi-spectral brain tissue segmentation using automatically trained k-Nearest-Neighbor classification. Neuroimage 2007; 37: 71–81.

23. Ikram MA, Vrooman HA, Vernooij MW, den Heijer T, Hofman A, Niessen WJ, et al. Brain tissue volumes in relation to cognitive function and risk of dementia. Neurobiol Aging 2010; 31:378–386.

24. Siepmann T, Boardman H, Bilderbeck A, Griffanti L, Kenworthy Y, Zwager C, et al. Long-term cerebral white and gray matter changes after preeclampsia. Neurology 2017; 88: 1256–1264.

25. Adank MC, Hussainali RF, Oosterveer LC, Ikram MA, Steegers EAP, Miller EC, Schalekamp-Timmermans S. Hypertensive disorders of pregnancy and cognitive impairment: A prospective cohort study. 2020; DOI:https://doi.org/10.1212/WNL.0000000000011363

26. Johnson AC, Cipolla MJ. The Cerebral Circulation During Pregnancy: Adapting to Preserve Normalcy. Physiology 2015; 30:139–147.

27. Fritz HC, Wittfeld K, Schmidt CO, Domin M, Grabe HJ, Hegenscheid K, et al. Current smoking and reduced gray matter volume - A voxel-based morphometry study. Neuropsychopharmacology 2014; 39: 2594–2600.

28. Arenaza-Urquijo EM, Landeau B, La Joie R, Mevel K, Mézenge F, Perrotin A, et al. Relationships between years of education and gray matter volume, metabolism and functional connectivity in healthy elders. Neuroimage 2013; 83: 450–457.

29. Molano JR. Hormone therapy and brain structure in postmenopausal women. Neurology 2016; 87: e100–e102.

30. Hammer ES, Cipolla MJ. Cerebrovascular Dysfunction in Preeclamptic Pregnancies. Curr Hypertens Rep 2015; 17: 1–13.

31. Soma-Pillay P, Suleman FE, Makin JD, Pattinson RC. Cerebral white matter lesions after pre-eclampsia. Pregnancy Hypertens 2017; 8: 15–20.

32. Logue OC, George EM, Bidwell GL. Preeclampsia and the brain: neural control of cardiovascular changes during pregnancy and neurological outcomes of preeclampsia. Clin Sci. (Lond) 2016; 130:1417–34.

33. Johannsen BMW, Larsen JT, Laursen TM, Bergink V, Meltzer-Brody S, Munk-Olsen T. All-cause mortality in women with severe postpartum psychiatric disorders. Am J Psychiatry 2016; 173: 635–642.

34. O’Hara MW, Wisner KL. Perinatal mental illness: Definition, description and aetiology. Best Pract Res Clin Obstet Gynaecol 2014; 28: 3–12.

35. Pawluski JL, Lonstein JS, Fleming AS. The Neurobiology of Postpartum Anxiety and Depression. Trends Neurosci. 2017; 40: 106–120.

36. Ziomkiewicz A, Sancilio A, Galbarczyk A, Klimek M, Jasienska G, Bribiescas RG. Evidence for the cost of reproduction in humans: High lifetime reproductive effort is associated with greater oxidative stress in post-menopausal women. PLoS One 2016; 11: e0145753.

37. Barha CK, Hanna CW, Salvante KG, Wilson SL, Robinson WP, Altman RM, Nepomnaschy PA. Number of children and telomere length in women: A prospective, longitudinal evaluation. PLoS One 12016;1:e0146424.

38. Pollack AZ, Rivers K, Ahrens KA. Parity associated with telomere length among US reproductive age women. Obstet Gynecol Surv 2018; 33: 736–744.

39. Kaptijn R, Thomese F, Liefbroer AC, Van Poppel F, Van Bodegom D, Westendorp RGJ. The trade-off between female fertility and longevity during the epidemiological transition in the Netherlands. PLoS One 2015; 10:e0144353.

40. Behl C, Widmann M, Trapp T, Holsboer F. 17-β estradiol protects neurons from oxidative stress-induced cell death in vitro. Biochem Biophys Res Commun 1995; 216:473–82.

41. Yim I, Tanner SL, Guardino C, Hahn-Holbrook J, Dunkel SC. Biological and psychosocial predictors of postpartum depression: systematic review and call for integration. Annu Rev Clin Psychol 2015; 11: 99–137.

42. Bridges RS. Long-term alterations in neural and endocrine processes induced by motherhood in mammals. Horm Behav 2016; 77:193–203.

43. Leuner B, Glasper ER, Gould E. Parenting and plasticity. Trends Neurosci 2010; 33: 465–473.

44. Salmaso N, Nadeau J, Woodside B. Steroid hormones and maternal experience interact to induce glial plasticity in the cingulate cortex. Eur J Neurosci 2009; 29: 786–794.

45. Kinder JM, Stelzer IA, Arck PC, Way SS. Immunological implications of pregnancy-induced microchimerism. Nat Rev Immunol 2017; 17: 483–494.

46. Zeng XX, Tan KH, Yeo A, Sasajala P, Tan X, Xiao ZC, et al. Pregnancy associated progenitor cells differentiate and mature into neurons in the maternal brain. Stem Cells Dev 2010; 19: 1819–1830.

47. Fjeldstad HE, Johnsen GM, Staff AC. Fetal microchimerism and implications for maternal health. Obstetric Medicine 2019; doi.org/10.1177/1753495X19884484.

48. Chan WFN, Gurnot C, Montine TJ, Sonnen JA, Guthrie KA, Nelson JL. Male microchimerism in the human female brain. PLoS One 20102; 7: e45592

49. Adams Waldorf KM, Nelson JL. Autoimmune disease during pregnancy and the microchimerism legacy of pregnancy. Immunol Invest 2008; 37: 631–644.

50. Cheng S.B, Davis S, Sharma S. Maternal-fetal cross talk through cell-free fetal DNA, telomere shortening, microchimerism, and inflammation. Am J Reprod Immunol 2018; 79: e12851.

51. Nassar D, Droitcourt C, Mathieu-d’Argent E, Kim MJ, Khosrotehrani K, Aractingi S. Fetal progenitor cells naturally transferred through pregnancy participate in inflammation and angiogenesis during wound healing. FASEB J 2012; 26: 149–157.

52. Gammill HS, Aydelotte TM, Guthrie KA, Nkwopara EC, Nelson JL. Cellular fetal microchimerism in preeclampsia. Hypertension 2013; 62: 1062–1067.

53. Marsland AL, Gianaros PJ, Kuan DCH, Sheu LK, Krajina K, Manuck SB. Brain morphology links systemic inflammation to cognitive function in midlife adults. Brain Behav Immun 2015; 48: 195–204.

54. Peltola MJ, Yrttiaho S, Puura K, Proverbio AM, Mononen N, Lehtimäki T, Leppänen JM. Motherhood and oxytocin receptor genetic variation are associated with selective changes in electrocortical responses to infant facial expressions. Emotion 2014; 14: 469–477.

55. Abraham E, Hendler T, Shapira-Lichter I, Kanat-Maymon Y, Zagoory-Sharon O, Feldman R. Father’s brain is sensitive to childcare experiences. Proc Natl Acad Sci 2014; 11: 9792–9797.

56. Parsons CE, Young KS, Petersen MV, Jegindoe Elmholdt EM, Vuust P, Stein A, Kringelbach ML. Duration of motherhood has incremental effects on mothers’ neural processing of infant vocal cues: a neuroimaging study of women. Sci Rep 2017; 7: 1–9.

57. Maupin AN, Hayes NJ, Mayes LC, Rutherford HJV. The application of electroencephalography to investigate the neural bases of parenting: a review. Parenting 2015; 15: 9–23.

58. Bick J, Dozier M, Bernard K, Grasso D, Simons R. Foster mother-infant bonding: associations between Foster mothers’ oxytocin production, electrophysiological brain activity, feelings of commitment, and caregiving quality. Child Development 2013; 84: 826–840.

59. Orchard ER, Ward PGD, Sforazzini F, Storey E, Egan GF, Jamadar, SD. Cortical changes associated with parenthood are present in late life. BioRxiv 2019; 0–3; https://doi.org/10.1101/589283

60. Roberts CL, Ford JB, Algert CS, Antonsen S, Chalmers J, Cnattingius S et al. Population-based trends in pregnancy hypertension and pre-eclampsia: an international comparative study. BMJ Open 2011; 1:e000101.

61. Eades CE, Cameron DM, Evans JMM. Prevalence of gestational diabetes mellitus in Europe: A meta-analysis. Diabetes research and clinical practice 2017; 129: 173–181.

62. Zwart JJ, Richters A, Ory F, de Vries JI, Bloemenkamp KW, van Roosmalen J. Eclampsia in the Netherlands. Obstetrics and gynecology 2008; 112: 820–827.

63. Bokslag A, Fons AB, Zeverijn LJ, Teunissen PW, de Groot CJM. Maternal recall of a history of early-onset preeclampsia, late-onset preeclampsia, or gestational hypertension: a validation study. Hypertension in pregnancy 2020; 39: 444–450.

